# A Nonsense Mutation in the PRKG2 Gene in Dalmatian Dogs with Chondrodysplasia

**DOI:** 10.1101/2025.03.18.644031

**Authors:** Mäkeläinen Suvi, Ekman Stina, Marjo K Hytönen, Lohi Hannes, Kyöstilä Kaisa, Simon Thomas, Andersson Göran, Hedhammar Åke, Hansson Kerstin, Bergström F. Tomas

## Abstract

Skeletal dysplasias encompass a diverse group of genetic disorders characterized by short stature and dwarfism. In humans, 771 types of skeletal dysplasia have been documented. Similar forms of these disorders have also been observed in dogs. The first cases of documented skeletal dysplasia in Dalmatian dogs were reported in the early 1980’s, with additional affected dogs observed in subsequent years. Careful radiological and histopathological examinations at the time revealed severe limb deformities, including shortened radii and ulnae, irregular growth plates and disrupted endochondral ossification. In this study, we applied whole-genome sequencing on samples collected in 1992 and identified a genetic variant in the *PRKG2* gene, introducing a premature stop codon (XM_038582312: c.T1601G, p.L534X). Genetic variants in *PRKG2* have previously been implicated in human acromesomelic dysplasia, a disorder affecting limb growth in young children. The *PRKG2*-encoded protein plays a crucial role in endochondral ossification, and if translated, the identified nonsense variant would result in a truncated protein lacking most of the catalytic domain. Extended screening of the genetic variant revealed its continued segregation in the current Dalmatian population. Furthermore, three recent cases of dwarfism in Dalmatians were found to be homozygous for the identified *PRKG2* nonsense variant. These findings provide compelling evidence for the role of *PRKG2* in Dalmatian dwarfism, resolving a decades-old genetic mystery in the breed.

## Introduction

Skeletal dysplasias represent a heterogeneous group of inherited diseases causing abnormalities in cartilage and bone, resulting in varying degrees of short stature and dwarfism(1). Collectively these disorders affect approximately 1 in 5000 people(2). The classification of different forms of the disorder have traditionally been based on radiographic and biochemical criteria. With the advances in molecular medicine and genomics, the nosology is increasingly being based on molecular criteria(3). The International Skeletal Dysplasia Society (ISDS) promotes research in the field and provides a comprehensive and updated list of different forms of inherited skeletal dysplasias (https://www.isds.ch). In the recent revision of inherited human skeletal dysplasias (Nosology of Genetic Skeletal Disorders: 2023 revision) describes 771 different types of skeletal disorders and 552 associated genes(3).

In domestic dogs, several forms of skeletal dysplasia have been described and for some breeds, dwarfism is part of the breed standard and a characteristic of the breed. Chondrodysplasia, a subset of skeletal dysplasias, primarily affects endochondral ossification – the process by which cartilage is converted into bone – leading to disproportionate limb shortening. This condition is the most commonly diagnosed form of skeletal dysplasia in dogs. To date, only a limited number of genetic factors underlying these disorders have been identified. Two *FGF4* retrogenes (*FGF4L1* on chromosome 18 and *FGF4L2* on chromosome 12) have been identified to cause dwarfism across many dog breeds(4–6). The dachshund is an example where the *FGF4L1* retrogene is fixed in the breed. Other genes include *PCYT1A* where a missense variant was proposed to cause disproportionate dwarfism in the vizsla breed(7) and a nonsense mutation in the *ITG10* gene likely result in a loss of protein function and cause chondrodysplasia in Norwegian Elkhound and Karelian Bear Dog breeds(8). The latter breed is also affected by pituitary from of dwarfism caused by an intronic variant in *POUF1F1* (9). A splice site mutation in the *PRKG2* gene was identified in the breed Dogo Argentino (10) and a partial deletion of the *SLC13A1* have been found in Miniature Poodles with a severe form of dwarfism(11). In addition, there are examples of syndromic diseases which result in disproportionate dwarfism. A form of oculoskeletal dysplasia, which is a hereditary syndrome comprising varying combinations of osteochondrodysplasia and ocular pathology. In Labrador retrievers the disease has been reported to be caused by a frameshift mutation in the gene *COL9A3* and in Samoyeds by a 1,267 bp deletion in the 5′ end of the *COL9A2* gene(12). A mild form of disproportionate dwarfism in Labrador retrievers have also been associated with a missense mutation in the *COL11A2* gene(13).

In Dalmatians, cases with severe chondrodysplasia were noted by breeders in the early 1980s. This was followed by sporadic cases in the following years and an autosomal recessive mode of inheritance was suspected. For one of the litters, born in 1992, a careful clinical examination including radiography was performed. Due to the severity of the condition, the affected dogs were euthanized and a necropsy was performed for three affected individuals. No molecular genetic study was initiated nor were the clinical findings published at the time and the cause of the disease remained unknown. However, blood samples, formalin embedded tissue samples, radiographic images as well as journal notes were archived. In this study, we revisited this cold case to investigate the genetic reason behind the skeletal dysplasia observed in Dalmatians from the 1990s and provide a thorough description of the clinical findings observed in these dogs.

## Results

### Clinical characterization indicates disproportionate dwarfism

In a litter of nine Dalmatians born in 1992, growth abnormalities were noted in four of the dogs at the age of two to three months. Three of the affected dogs as well as one unaffected littermate were clinically examined and radiographed at the age of four months (**Fig 1**). A video recording from 1992 shows the affected female at the age of 1.5, 3 and 4 months, as well as the male affected dog and the unaffected female littermate at the age of 4 months (**S1 Video**). All three affected dogs had short legs and showed clear gait abnormalities. The front legs were curved with an outward-angled elbow joint and an outward rotation of the paw. Due to the severity of the condition, the affected dogs were euthanized and a necropsy was performed. Among the affected dogs, the radiological examination revealed angular limb deformities. Both the ulna and the radius were found to be short and wide with a more pronounced shortening of the ulna and a cranially curved radius (**Fig 1B and 1D**). In the elbow, an intraarticular stairstep formation and a proximally displaced ununited anconeal process were observed (**Fig 1D**). The asynchronous growth of the elbow joints was confirmed by necropsy (**Fig 1E**). The distal ulna appeared wider with an irregular growth plate. Microscopically, the growth plates of the distal ulna and radius were irregular showing disorganized areas of resting, proliferative and hypertrophic zones (**Fig 2A-C**). Areas of chondronecrosis and eosinophilic streaks of remnants of vessels were found in the resting/proliferative zones (**Fig 2B and 2C**). The regular pattern with different growth regions were obliterated and groups of proliferative and hypertrophic chondrocytes were intermingled (**Fig 2A-D**). The growth plates were in many areas also found to be thinner than normal (**Fig 2C**). Areas of normal growth plate differentiation could be identified in distal radius (**Fig 2D**).

**Fig 1.**
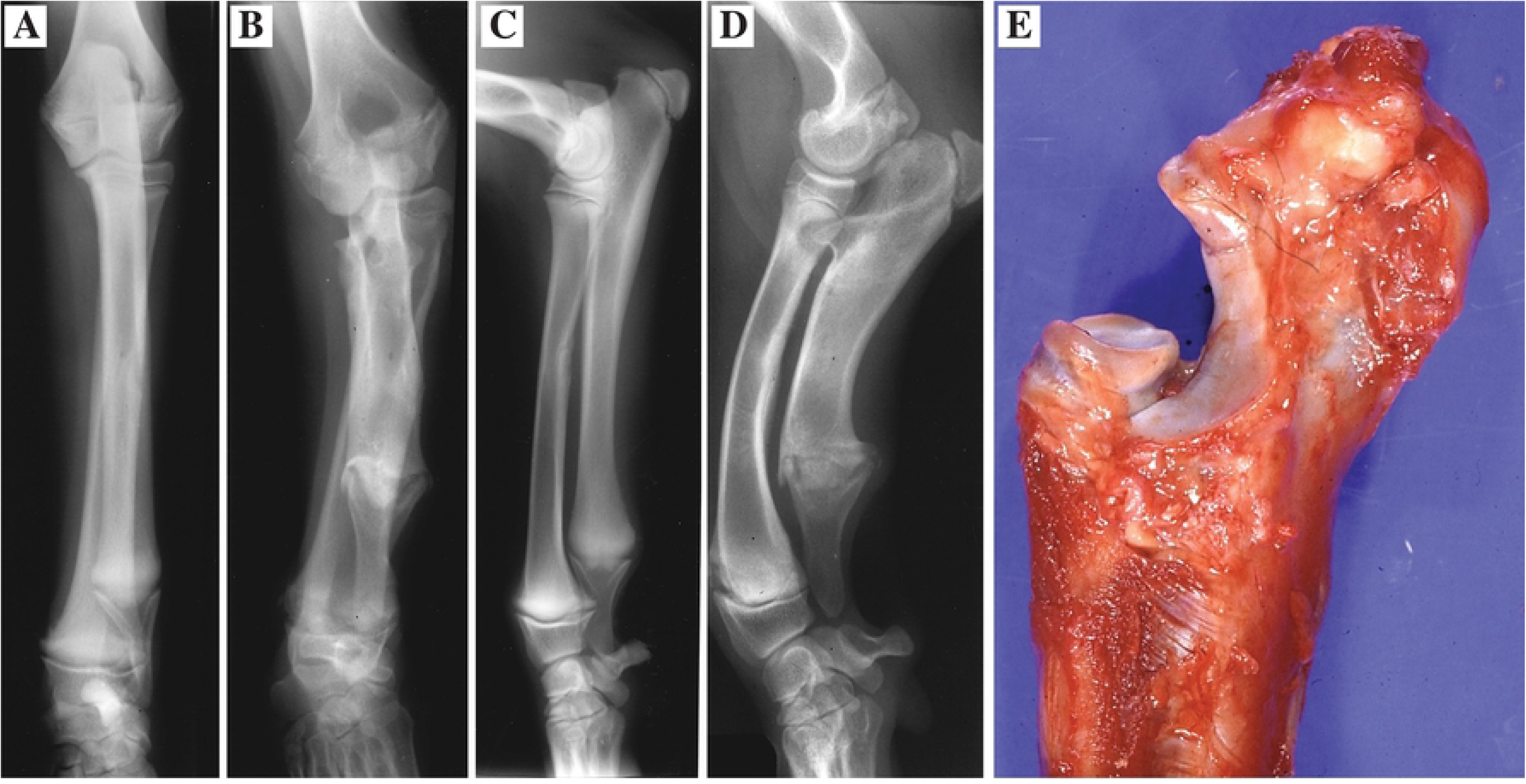
Diagnostic imaging and macroscopic examination. Craniocaudal projections of the antebrachium of a **(A)** non-affected and an **(B)** affected dog as well as mediolateral projections of a **(C)** non-affected and an **(D)** affected dog at the age of 4 months. In the affected dog **(B and D**) the most obvious changes are a pronounced shortening of the ulna and cranial curving of the radius with a difference in length between the two bones giving rise to an intra-articular stairstep formation. **(E)** A step was developed in the elbow joint with shorter ulna in relation to radius.

**Fig 2.**
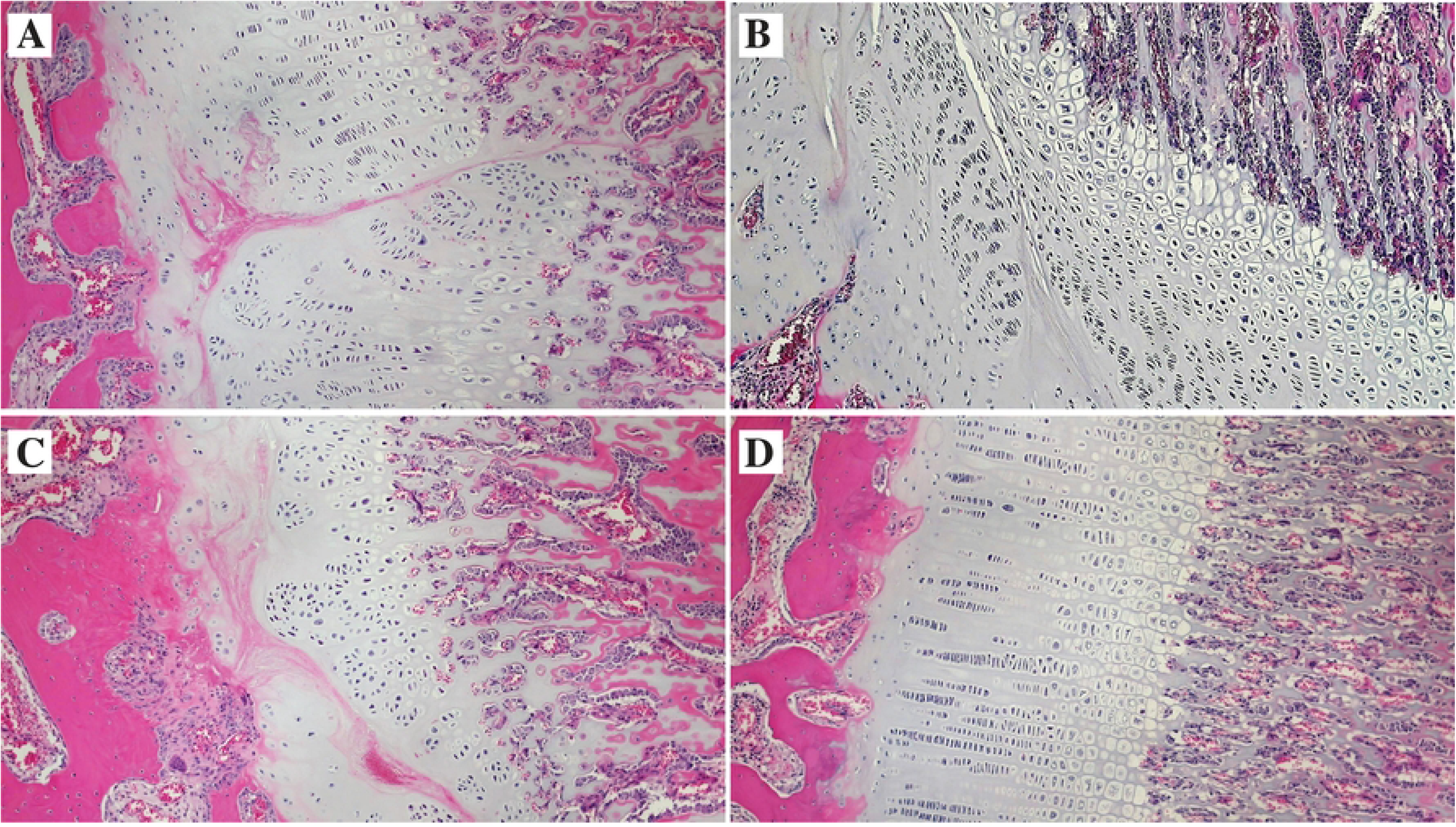
Histopathology. Light microscopic examination of the epiphysal plates, stained with hematoxylin and eosin, from affected dogs at the age of four months (**A)** Irregular proliferative and resting zones with areas of chondronecrosis including eosinophilic streaks of remnants of vessels present in the radial physis. **(B)** Areas of chondronecrosis in the resting and proliferative zones of the ulnar physis. **(C)** Irregular proliferative and resting zones with areas of chondronecrosis, including eosinophilic streaks of remnants of vessels seen in a thin radial physis **(D)** Regular differentiation of resting, proliferative and hypertrophic zones in an area of the radial physis.

Radiological examination of the hindlegs were performed on the affected male dog and one of the affected female dogs. The results showed that the tibia and fibula were shorter than normal. Irregular chondro-osseous junctions between growth cartilage and bone of the metaphyses and epiphyses in humerus, femur and tibia were present in the female dog. Chondronecrotic areas of these growth plates were also found. Most of the growth plates from femur, tibia, humerus and radius of the affected male dog were characterized by morphologically normal growth cartilage with an active endochondral ossification.

The axial skeleton of the affected dogs was shorter in length compared to the unaffected dogs. The thoracic disc (Th12-Th13) had an outer wall of dense connective tissue with an inner layer of laminar fibrocartilage. The nucleus pulposus consisted of rounded cells in an amorphous viscous fluid. There was no evidence of a hyaline-type of cartilage in the inner perinuclear layer, as is described in achondroplastic and chondrodystophic dogs. The microscopic appearance of the growth cartilage in the ribs was normal with an active endochondral ossification.

### Whole genome sequencing identifies a candidate variant in *PRKG2*

We sequenced the genomes of two affected males as well as the unaffected parents at an average sequencing depth of 18.6X. Conditional filtering, assuming an autosomal recessive mode of inheritance, led to the identification of 106 non-synonymous SNVs and three nonsense variants. To narrow down the number of candidate variants further we then filtered against known variants found in a dataset of 1,987 dogs(14). As a result, five variants remained, comprising four non-synonymous substitutions and one nonsense (stop-gain) variant (**S1 Table**).

We utilized *in silico* tools to predict the functional impact of the identified nonsynonymous substitutions. Variants in three genes – olfactory receptor family 52 subfamily A member 18 (*OR52A18*), NLR family pyrin domain containing 14 (*NLRP14*) and olfactory receptor family 52 subfamily A member 5C (*OR52A5C*) – were predicted to be tolerated and benign. The functional effect of the variant in the gene *LOC119876480* (translation initiation factor IF-2-like), could not be predicted. The gene belongs to a predicted subset of NCBI RefSeq genes, annotated *ab initio* by an automated computational analysis, and has no RefSeq counterparts in other species. Moreover, the function of the four genes with nonsynonymous substitutions indicated no association to skeletal growth. Consequently, based on the protein function, known disease associations, and predicted effect of the mutation, none of the non-synonymous substitutions were regarded as likely candidates for the condition (**S1 Table**).

The nonsense variant (canFam4 chr32:34,701,558T>G, XM_038582312: c.T1601G, p.L534X) was located in the protein kinase cGMP-dependent 2 (*PRKG2*) gene. This gene is among the 552 genes implicated in human skeletal dysplasias (Nosology of Genetic Skeletal Disorders: 2023 revision)(3). Moreover, a splice site variant in the same gene has previously been implicated in canine disproportionate dwarfism in the Dogo Argentino breed (10). The nonsense mutation was thus considered a strong candidate variant for the Dalmatian disorder. Two different *PRKG2* transcripts are described to date, producing 762 and 733 amino acid residues long protein products, respectively. Both isoforms include a ligand binding domain and a serine/threonine kinase catalytic domain. The identified nonsense variant was predicted to truncate the highly conserved catalytic domain of the protein, resulting in a lysine to premature stop change at amino acid residues 534 or 505, of the respective transcripts. The 762 amino acid residues long human (UniProtKB accession number Q13237-1) and canine (UniProtKB A0A8C0NE14) protein products exhibit 96.7% sequence identity over their entire length (**S1 Fig**) and the lysine at amino acid position 534 is highly conserved among mammals.

### Genetic validation supports the causality of the nonsense variant in *PRKG2*

Next, we validated the inferred *PRKG2* genotype in the four whole-genome sequenced dogs and the remaining littermates, including one affected female as well as four unaffected siblings (**S2 Fig**). All three affected dogs were homozygous for the *PRKG2* nonsense variant. The clinical examination also included a second clinically affected female dog (**Fig 3A**), but no blood sample had been collected at the time. Attempts to extract DNA from paraffin-embedded tissue of this dog were not successful and thus, we could not retrieve the genotype. Of the four unaffected individuals, two dogs were wildtype, including the unaffected female that’s was subjected to radiographic examination, and one dog was heterozygous. Surprisingly, the fourth littermate, assumed at the time to be unaffected, was tested homozygous for the genetic variant. We then contacted the owner of this fourth dog, who confirmed that the dog was indeed considered healthy and passed away at the age of 16 years. However, according to the owner, the dog was 8 cm shorter in height at withers compared to the owner’s second, unaffected Dalmatian female and the physical appearance of the dog was sturdier than normal.

**Fig 3.**
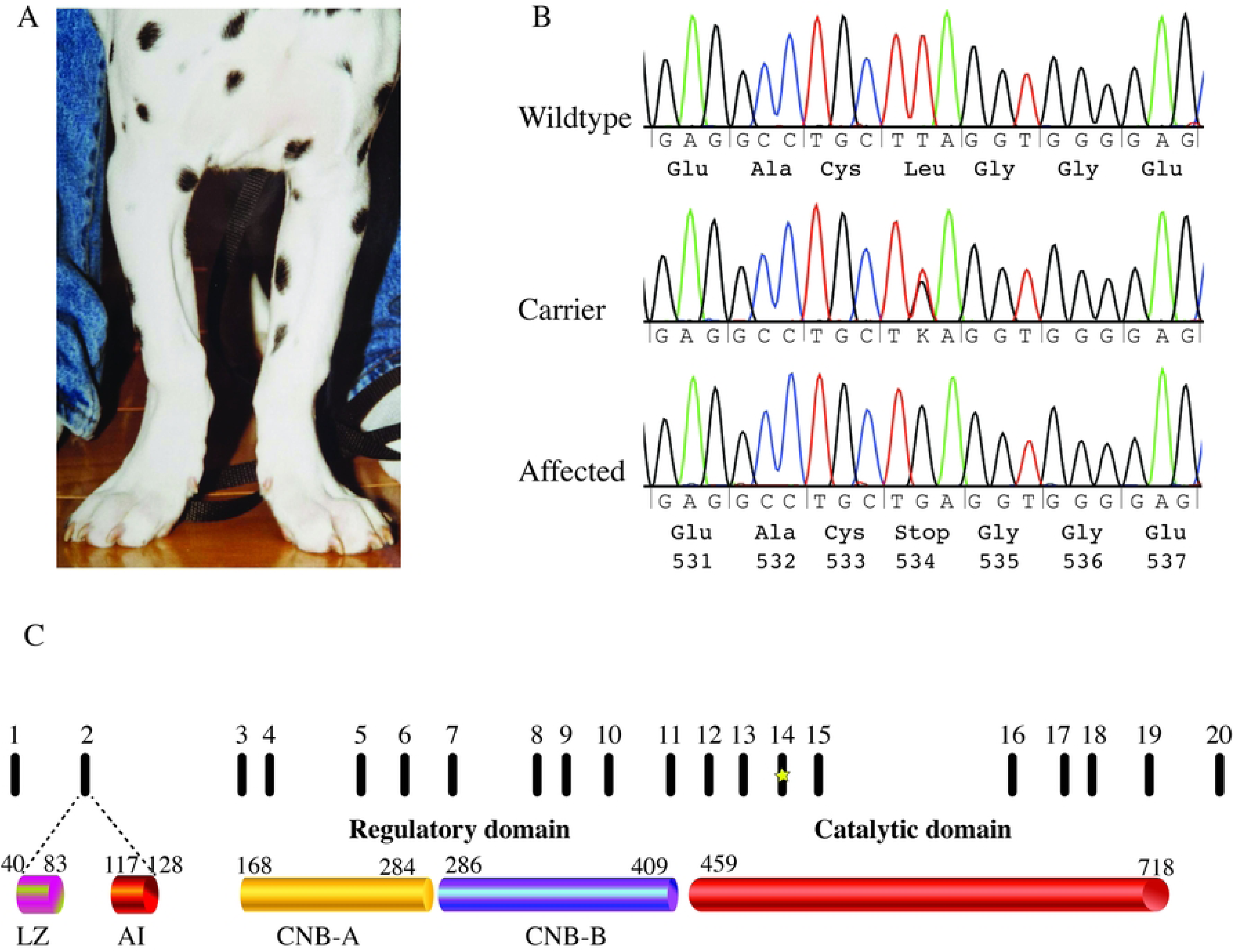
Nonsense mutation in the canine PRKG2 gene. **(A)** Affected female at approximately two months of age **(B)** Sanger sequencing traces spanning positions chr32:34,701,547-34,701,568 (canFam4) in exon 14 of the PRKG2 gene. The identified T to G change at position 34,701,558 affects the second position of codon 534 and results in premature stop codon (p.L534X). **(C)** Graphic representation of the exons in the PRKG2 gene and the encoded protein domains (LZ = leucine zippers, Al = autoinhibitory domain, CNB = cyclic nucleotide-binding) including the catalytic domain (aa 459-718). The premature stop codon at amino acid position 534 is indicated in exon 14 with a star.

To further evaluate the concordance of the nonsense variant with the disease phenotype, we genotyped an additional 291 Dalmatians using Sanger sequencing (**Fig 3B**) or real-time PCR. Ten of these samples were collected in the 1990s. The remaining samples were obtained more recently and included three dogs diagnosed with chondrodysplasia within the past decade. Notably, all three affected dogs were homozygous for the candidate variant. Among the remaining 288 unaffected individuals, 21 were heterozygous for the variant, while the other 267 were homozygous for the wild-type allele.

## Discussion

Family based whole-genome sequencing has become a powerful strategy to identify disease causing variants for relatively rare diseases with autosomal recessive mode of inheritance. In canine genetics, this was first shown in 2016, when a missense variant in the gene family with sequence similarity 83 member G (*FAM83G*) was identified as a likely candidate for hereditary footpad hyperkeratosis in Kromfohrländer dogs(15). In the current study, we used the same approach to identify the genetic cause of a skeletal dysplasia in a family of Dalmatian dogs.

Whole-genome sequencing identified a nonsense variant (canFam4 chr32:34,701,558T>G, c.T1601G, p.L534X) in the *PRKG2* gene which was considered as a strong candidate for the condition in the Dalmatian dogs. The *PRKG2* gene, encoding the cGMP-dependent protein kinase 2 (cGKII), plays a critical role in regulating endochondral ossification and bone growth. This was initially demonstrated in *Prkg2*-deficient mice(16). The mice had normal embryonic and fetal development but after birth, their growth was abnormal and dwarfism and short limbs became apparent. This is similar to the Dalmatian dogs in this study, where according to the breeders, no obvious signs of skeletal dysplasia were observed during the first weeks after birth.

The *Prkg2*-deficient mice showed normal development of membranous bones in the cranial vault, suggesting that the effect was specific to endochondral ossification. Most bones in the skeleton, including appendicular bones (*e.g.* femur and tibia), vertebrae and medial clavicles, are formed by endochondral ossification, involving chondrogenesis from mesenchymal stem cells followed by chondrocyte differentiation to terminal hypertrophic chondrocytes. During later stages of chondrogenesis, in the process of hypertrophic differentiation, chondrocytes enlarge, terminally differentiate, mineralize, and ultimately undergo apoptosis (for review see: (17–19). cGKII, encoded by *PRKG2,* is critical for the process of hypertrophic differentiation. The protein kinase is activated by the increase of cyclic guanosine monophosphate (cGMP) levels as a result of C-type natriuretic peptide (CNP) stimulation of its receptor natriuretic peptide receptor-B (NPR-B) (20, 21). The activated cGKII inhibits the mitogen-activated protein kinase (MAPK) cascade at the level of RAF-1 and may also regulate and phosphorylate the transcription factor SOX9, resulting in hypertrophic chondrocyte differentiation (**Fig 4**) (22, 23).

**Fig 4.**
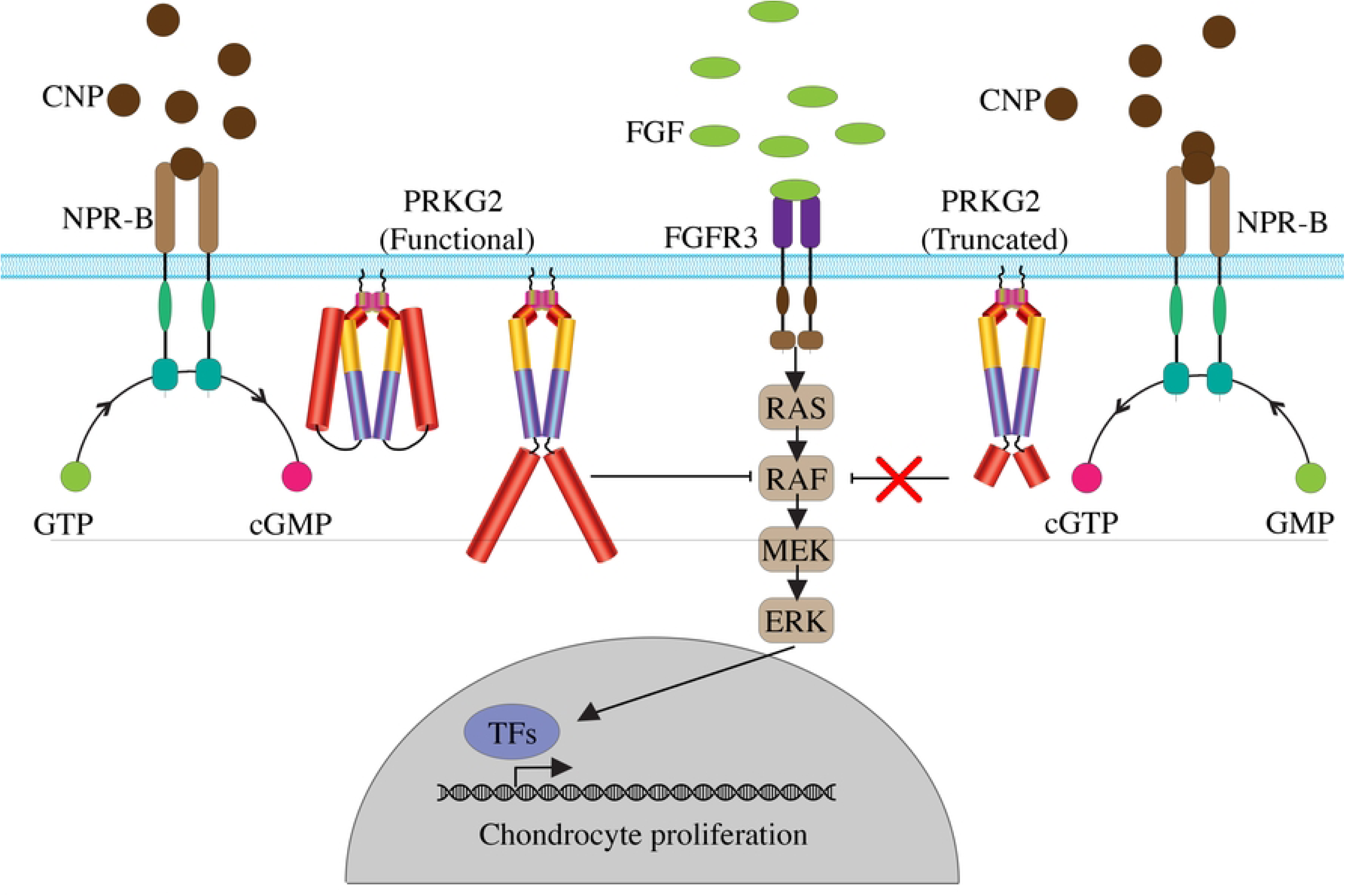
PRKG2 (cGKII) in CNP-mediated endochondral ossification illustrating the functional (left) and a hypothetical truncated PRKG2 (right). Upon C-type natriuretic peptide (CNP) binding to natriuretic peptide receptor-B (NPR-B), intracellular cyclic guanosine monophosphate (cGMP) levels increase, and act as a second messenger by the activation of cGMP-dependent protein kinase 2 (cGKII/PRKG2) homodimer. Activated cGKII inhibits the MAPK cascade (FGFR3/RAS/RAF/MEK/ERK) at the level of RAF, thus antagonizing fibroblast growth factor (FGF)-induced MAPK signaling and thereby promoting hypertrophic chondrocyte differentiation. The nonsense mutation identified in this study introduces a premature stop codon at amino acid position 534 within the catalytic domain of cGKII, illustrated by the shortened red domains in the homodimer, likely resulting in a loss of catalytic activity. This loss is predicted to impair the inhibition of MAPK signaling, leading to disruption of the normal switch from proliferating chondrocytes to hypertrophic differentiation. TFs = transcription factors.

The canonical transcript of *PRKG2* in both human and dogs encodes for a protein with 762 amino acids which forms a homodimer linked by leucine zippers at the N-terminus(24, 25). The regulatory ligand binding domain consists of the two cyclic nucleotide-binding sites (CNB-A and CNB-B) with different affinities for cGMP binding (25, 26) (**Fig 3C**). The catalytic domain spans over amino acid residues 459-718. In the absence of cGMP, the catalytic domain interacts with the autoinhibitory regulatory domain. Increased cellular levels of cGMP binding to the regulatory region, results in a conformational change of cGKII and allows the homodimer to phosphorylate the substrate which in hypertrophic differentiation is RAF-1(22, 23, 25, 26). The identified nonsense variant in this study results in a premature stop codon early in the catalytic domain, amino acid position 534. If translated, we would expect lack of catalytic activity, and an inability of cGKII to inhibit MAPK resulting in dysregulation of chondrocyte differentiation (**Fig 4**).

Five disease-causing *PRKG2* alleles (27, 28) have been identified in humans to date. These variants cause PRKG2-related acromesomelic dysplasia (acromesomelic dysplasia 4, AMD4, MIM 619636) and possibly PRKG2-related spondylometaphyseal dysplasia (Pagnamenta type (SMDP, MIM 619638) (29). Acromesomelic dysplasia results in shortened forearms and forelegs (mesomelia) and abnormal shortening of the bones in the hands and feet (acromelia). The clinical findings in the affected dogs, including shortening and bending of the long bones align with the mesomelic changes in the affected humans, but unlike the human patients, the affected dogs showed no marked shortening of the distal appendicular bones.

Evidence suggests that all five identified human variants (a nonsense (p.Arg569*), a splice-site (c.1635-1 G>A), two frameshift variants (p.Asn164Lysfs*2 and p.Asp761Glufs*34) and a missense variant (p.Val470Gly) cause loss of protein function (**S1 Fig**). In the dog breed Dogo Argentino, a splice donor variant in the *PRKG2* gene (PRKG2:XM_022413533.1, c.1634+1G>T), has been implicated in a disproportionate form of dwarfism (10). Similar to the identified candidate variant in the Dalmatian dogs and the human nonsense variant (p.Arg569*), the splice donor variant identified in Dogo Argentino(10) is also located in the mRNA corresponding to the catalytic domain of the encoded protein . Furthermore, naturally occurring mutations in the *PRKG2* gene have been shown to cause dwarfism in rats (Komeda miniature rat Ishikawa, KMI) (22) and American Angus cattle (30).

This study comprised seven clinically affected dogs. Blood samples were available for six of these individuals, all of which were found to be homozygous for the identified nonsense variant in the *PRKG2* gene. Notably, genotyping of one female littermate unexpectedly revealed a homozygous genotype for the nonsense variant. To assess the possibility of sample misidentification, we attempted to extract genomic DNA from a formalin-fixed, paraffin-embedded (FFPE) tissue sample of the seventh clinically affected female, for whom no blood sample was available, with the aim of comparing the DNA profile of this affected dog and the blood sample from the homozygous female littermate. However, DNA extraction was unsuccessful, likely due to prior decalcification treatment of the tissue during fixation and paraffin embedding. Based on the owner’s observations, the homozygous littermate may have exhibited mild growth retardation. The finding of the mildly affected homozygous individual suggest that homozygosity for the identified nonsense variant in *PRKG2* may result in considerable phenotypic variation.

To further assess the variant’s association with the condition, we genotyped additional Dalmatian dogs sampled during the 1990s as well as samples collected recently, but did not identify other unaffected dogs homozygous for the variant. Based on these findings, we conclude that the identified variant is a likely causative genetic variant for the disorder. Furthermore, identification of this nonsense variant allele in the recent samples from the Dalmatian population indicates that the variant remains segregating within the breed. These results highlight the importance of considering this variant in future breeding strategies to mitigate the occurrence of *PRKG2*-related chondrodysplasia in Dalmatians.

## Material and methods

### Animals and samples

Samples from the 1992 family of Dalmatians were used in the initial analysis of this study. In addition to the samples from dam and sire, we had access to blood samples from three affected individuals (two males and one female), as well as four unaffected siblings. The litter also included one affected female from which we only had access to radiographs and FFPE tissue, as well as one unaffected male which was not sampled. We also had access to ten additional whole blood samples from Dalmatians collected during the 1990’s as well as 281 samples collected more recently. All dogs included in this study were privately owned Dalmatians. The samples were obtained with informed dog owners’ consent. Ethical approval was granted by the regional animal ethics committees (Uppsala djurförsöksetiska nämnd; Dnr C12/15, Dnr 5.8.18-15533/2018, and Dnr 5.8.18-04682/2020 and the Animal Ethics Committee of State Provincial Office of Southern Finland; ESAVI/25696/2020).

Genomic DNA was extracted using 0.2-1 ml blood on a QIAsymphony SP instrument and the QIAsymphony DSP DNA Kit (Qiagen, Hilden, Germany) or by using a semi-automated chemagic 360 instrument (PerkinElmer Chemagen Technologie GmbH). We also attempted to extract DNA from a FFPE tissue sample (ribs) of the second clinically affected female in the 1992-litter using a truXTRAC® FFPE total NA Ultra Kit (Covaris Inc., Woburn, MA). However, the extraction did not yield any DNA, likely due to the decalcification process described above.

Three affected littermates (one male and two females) were necropsied two hours after euthanasia at an age of four months. A routine necropsy was performed for all three dogs, including a macroscopic and microscopic examination of the distal and proximal ulna, radius, tibia, proximal femur and humerus, and thoracic vertebrae 12-13, including the disc.

### Diagnostic imaging

Three affected and one unaffected female dog from the litter were radiographed at the age of four months. In the unaffected and two of the affected dogs, radiographs were taken of the axial skeleton in left lateral recumbency. In all three affected dogs, antebrachium of both front limbs, and tibia and fibula of both hind limbs were radiographed in mediolateral and craniocaudal projections.

### Histopathological examination

The specimens were immersed in an aqueous solution of 4% buffered formaldehyde, decalcified in citrate-buffered formic acid, embedded in paraffin and cut into approximately 6 μm and stained with hematoxylin and eosin (H&E).

### Whole-genome sequencing

Genomic DNA from sire, dam, and two affected male offspring from the 1992 litter was fragmented using the Covaris M220 instrument (Covaris Inc., Woburn, MA), according to the manufacturer’s instructions. Libraries with insert sizes of 550 bp were constructed following TruSeq DNA PCR-Free Library Prep protocol. The libraries were multiplexed and sequenced on a NextSeq500 instrument (Illumina, San Diego, CA) for 100 x 2 and 150 x 2 cycles using the High Output Kit and High Output Kit v2, respectively. The raw base calls were de-multiplexed and converted to fastq files using bcl2fastq v.2.15.0 (Illumina).

The data from the sequencing run was trimmed for adapters and low-quality bases using Trimmomatic v.0.32(31), and aligned to the canine reference genome CanFam4 UU_Cfam_GSD_1.0(32), using Burrows-Wheeler Aligner (BWA) (33)(bwa-mem2 v.2.2.1-20211213-edc703f)(34). Aligned reads were sorted and indexed using Samtools v.1.16 (35) and duplicates were marked using Picard v.2.27. 5(36). The BAM files were realigned and recalibrated with GATK v.4.3.0(37). SNPs and INDELs were called for each sample separately, after which the genomic variant calls were merged and genotyped across the four samples simultaneously using standard hard filtering parameters according to GATK Best Practices (38) recommendations. Variants annotated in the exonic regions with ANNOVAR v.2020.06.08 (39) using NCBI *Canis lupus familiaris* Annotation Release 106 were selected for further evaluation. The sequence data were submitted to the European Nucleotide Archive (ENA) with the accession number PRJEB79353.

We then used conditional filtering to filter for variants with autosomal recessive inheritance pattern, expecting the affected individuals to be homozygous for the variant and the sire and dam to be heterozygous for the variant. The identified variants were further filtered against known variation in the publicly available data from Dog10k consortium (https://zenodo.org/records/8084059)(14).

To predict the effects of amino acid changes on protein function, we evaluated the identified missense variants using PolyPhen-2 v2.2.3r406 (40) and SIFT Sequence for Single Protein Tools (41). Moreover, we used the human gene database GeneCards (www.genecards.org) to retrieve NCBI gene summary to predict the function and disease-association for each of the genes with candidate variants (42, 43). The pairwise sequence alignment (EMBOSS Needle) was made using UniProtKB fasta files with accession number Q13237-1 (human) and A0A8C0NE14 (canine).

### Genotyping

We designed primers using Primer3 v.0.4.0 (44, 45) (**S2 Table**) with and without m13 tails to amplify the variant canFam4 chr32:34,701,558T>G (c.T1601G:p.L534X) in *PRKG2* gene. The Sanger sequencing reaction was either performed using the BigDye™ Direct Cycle Sequencing Kit with the reactions loaded on Applied Biosystems 3500XL Series Genetic Analyzer system (Applied Biosystems, Thermo Fisher Scientific, Waltham, MA) or by amplification without m13 tails after which the reactions were treated with exonuclease I (New England Biolabs, Ipswich, MA) and rapid alkaline phosphatase (Roche Diagnostics, Solna, Sweden) before being loaded on an ABI 3730 capillary sequencer (Applied Biosystems, Thermo Fisher Scientific, Waltham, MA). The sequences were analyzed using CodonCode Aligner (CodonCode Corporation, Dedham, MA) and Unipro UGENE v1.32.0 software (46–48). The variant was also genotyped with a custom made TaqMan Real-Time PCR Assay (for primers and probes see (**S2 Table**). The amplifications were performed in a 10μl volume with the TaqPath™ ProAmp™ Master Mix amplification and detection was made on a StepOnePlus instrument (Applied Biosystems, Thermo Fisher Scientific).

## Acknowledgements

We would like to thank Ulla-Britt, Kenneth and Mikaela Lodelius (Kennel Courbettes) for providing detailed information about the 1992 litter as well as photos and video recordings. We would also like to thank Inger Hagbohm (Kennel Boing) for providing a detailed historical information about skeletal dysplasia in the breed and Erna Kuipers (Kennel Namaras) for coordinating the collection of samples and providing information about the recent cases in Europe. Moreover, we are grateful to the Finnish Dalmatian Club (Suvi Lukjanov and Erica Nyman) for their contributions and to all the owners and breeders of Dalmatians who have contributed to the research We would also like to thank Sini Karjalainen for technical assistance, Susanne Gustafsson at the SLU canine biobank for preparing and extracting samples, Tytti Vanhala for genotyping and Daniela Hahn for sequencing library preparation. The whole-genome sequencing analysis was enabled by resources provided by the National Academic Infrastructure for Supercomputing in Sweden (NAISS), partially funded by the Swedish Research Council through grant agreement no. 2022-06725.

## Supporting information captions

**S1 Video**. **Video recording from 1992.** The footage shows an affected female at 1.5, 3, and 4 months of age, along with her affected male littermate and an unaffected female littermate at 4 months of age. Notice the difference in front limb conformation between the affected dogs and the unaffected littermate at 4 months of age and the dramatic development of angular limb deformation of the front legs in the affected female between 1.5 and 3 months.

**S1 Fig**. **Protein sequence alignment.** The pairwise alignment (Needle) of the human (UniProtKB accession number Q13237-1) and canine (UniProtKB A0A8C0NE14) 762 amino acid long PRKG2 protein show 96.7% (735/762) identity. The identified nonsense variant at position 534 in this study as well as the previously described splice donor variant in Dogo Argentino is indicated on the canine sequence as well as disease-causing variants in human patients.

**S2 Fig**. **Pedigree and genotypes of the Dalmatian litter from 1992.** Pedigree showing the unaffected parents and nine offspring. Offspring 1 was clinically affected and included in the clinical study. Offspring 2 was clinically affected and included in the clinical study, but no blood sample was available for this dog. Offspring 3 was not clinically examined and assumed healthy. According to the owner this dog was small and sturdier than normal. Offspring 4 was healthy and clinically examined at the age of four months. Offspring 5 and 6 were assumed healthy and were not clinically examined. Offspring 7 was assumed healthy and no blood sample was available for this dog. Offspring 8 and 9 were clinically affected and whole-genome sequenced. Offspring 9 was included in the clinical study. Circles represent females, squares represent males. G/G = homozygous for the *PRKG2* nonsense mutation. T/G = heterozygous for the *PRKG2* nonsense mutation. T/T homozygous wildtype.

**S1 Table**. **List of candidate variants.** Coding sequence variants identified as private for the Dalmatian family, including the predicted effect of the variants based on SIFT and Polyphen2.

**S2 Table**. **Primers and probes used in the analysis.**

